# Oiling the gears of memory: quercetin exposure during memory formation, consolidation, and recall enhances memory in *Lymnaea stagnalis*

**DOI:** 10.1101/2021.06.24.449824

**Authors:** Veronica Rivi, Anuradha Batabyal, Cristina Benatti, Johanna MC Blom, Fabio Tascedda, Ken Lukowiak

## Abstract

Memory formation (short-term, intermediate-term, and long-term) is an integral process of cognition which allows individuals to retain important information and is influenced by various intrinsic and extrinsic factors. A major extrinsic factor influencing cognition across taxa is diet, which may contain rich sources of molecular agents with antioxidant, anti-inflammatory, and memory enhancing properties that potentially enhance cognitive ability. A common and abundant flavonoid present in numerous food substances is quercetin (Q) which is also known to upregulate cyclic AMP response element binding protein (CREB) in several animals including our model system *Lymnaea stagnalis*. Since CREB is known to be involved in long term memory (LTM) formation, we investigated the role of Q-exposure on memory formation, consolidation, and recall during operant conditioning of aerial respiratory behaviour in *Lymnaea*. Snails were exposed to Q 3h before or after training to ascertain its effects on LTM. Additionally, we investigated the effect of the combined presentation of a single reinforcing stimulus (at 24h post-training or 24h before training) and Q-exposure on both LTM formation and reconsolidation. Our data indicate that Q-exposure acts on the different phases of memory formation, consolidation, and recall leading to enhanced LTM formation.

**Summary Statement:** Quercetin enhances long-term memory formation acting on the different phases of memory formation, consolidation, and recall.

## Introduction

Learning is the ability to acquire new knowledge, whereas memory is the process of storing, recalling, and reconstructing that knowledge over time (Kandel et al., 2014; Zlotnik and Vansintjan, 2019). Memories are not only essential for survival but are also major determinants of “*who we are’’* (Milner et al., 1998). In fact, by regulating decision-making processes and influencing emotional reactions, they shape our very identity (Brem et al., 2013; Wilson and Ross, 2003).

Memory, based on the duration of its persistence, can be classified into short-term memory (STM, lasting minutes), intermediate-term memory (ITM, lasting 2–3 h), and long-term memory (LTM, persisting for at least 24h and sometimes for the entire lifetime; Braun and Lukowiak, 2011; Lukowiak et al., 2000; Sangha et al., 2003a,b). LTM is functionally dynamic and includes several temporal phases, such as encoding, consolidation, retrieval, and reconsolidation (Bisaz et al., 2014; Marra et al., 2013; Lukowiak et al., 2000; Rosenzweig et al., 1993). In recent decades, extraordinary progress has been made in demonstrating the molecular and cellular mechanisms underlying LTM, providing insight into how memories might also be enhanced. In this regard, studies in invertebrate models such as *Aplysia californica* (Conte et al., 2017; Hawkins et al., 2006; Philips et al., 2006) and *Drosophila melanogaster* (Davis, 2011; Isabel et al., 2004; Matsuno et al., 2015; Widmer et al., 2018) have paved the way to the elucidation of the conserved molecular and cellular basis of memory across *taxa*. A key feature of LTM is that a newly encoded memory initially exists in a fragile state and, over time, becomes increasingly resistant to disruption. With retrieval or reactivation, a consolidated memory can re-enter a labile state until it is restabilized through a process of reconsolidation (Nader et al., 2000; Alberini, 2009, 2011; Bisaz et al., 2014; Squire et al., 2015). During both consolidation and reconsolidation processes, memory is susceptible to modification by environmental factors, such as stress and lifestyle choices (McKenzie and Eichenbaum, 2011).

One key lifestyle choice that impacts the strengthening or weakening of the memory trace is diet (Swinton et al., 2018). Growing evidence suggests that a group of dietary-derived phytochemicals, known as flavonoids, may improve LTM formation, consolidation, storage, and retrieval (Ayaz et al., 2019; Fruson et al., 2012; Haleagrahara et al., 2011; Havsteen, 2002; Knezevic and Lukowiak, 2014; Letenneur et al., 2007; Socci et al., 2017). A multitude of pharmacological studies reported that regular dietary consumption of flavonoids and flavonoid-rich foods can improve various cognitive dysfunctions and dementia-like alterations in different animal models (Bakoyiannis et al., 2019; Macready et al., 2009; Mastroiacovo et al., 2015; Socci et al., 2017; Spencer, 2007a; Yevchak et al., 2008). The positive cognitive effects of flavonoids in mammals have been attributed to the protection of neural functioning, stimulation of neuronal regeneration, and increased blood flow to the brain (Macready et al., 2009; Mastroiacovo et al., 2015; Schroeter et al., 2007; Spencer, 2007; Vauzour et al., 2007; Williams and Spencer, 2012; Yevchak et al., 2008).

Quercetin (Q, 3,3′,4′,5,7◻pentahydroxyflavone) is one such flavonoid which is widely distributed in fruits and vegetables, like apples, berries, onions, asparagus, capers, and red leaf lettuce (Manach et al., 2004, 2005). Q has antioxidant properties and is able to scavenge free radicals and protect neuronal cells from neurotoxicity caused by oxidative stress (Ahmad et al., 2015; Cushnie and Lamb, 2005; Eggler et al., 2008; Kong et al., 2000; Morel et al., 1993a; Robak and Gryglewski, 1988). Unfortunately, to date, the vast majority of the studies investigating the effects of Q on memory has been performed on aged individuals or animal models of neurodegenerative diseases (Ansari et al., 2009; Bhutada et al., 2010; Broman-Fulks et al., 2012; Haleagrahara et al., 2011; Jayasena et al., 2013; Karimipour et al., 2019; Khan et al., 2018; Mandel and Youdim, 2004; Morel et al., 1993; Nakagawa et al., 2016; Reznichenko et al., 2006; Spencer, 2007, 2008; Williams and Spencer, 2012; Yoshida et al., 1990). Thus, there is a paucity of studies about the memory-enhancing effect of Q on healthy individuals. However, studies involving Q treatment on cognitively impaired animal models showed several molecular mechanisms that can enhance memory. Among them, the extracellular signal-regulated kinase (ERK1/2) and the protein kinase B (PKB/Akt) signalling pathways are important as they activate the response element-binding protein (CREB) family of transcription factors which, in turn, play a key role in regulating the genes necessary for synaptic plasticity (Finkbeiner et al., 1997; Spencer, 2007; Spencer et al., 2003; Vauzour et al., 2007). In particular, the transcriptional activator CREB1 is a core component of the molecular switch that converts STM to LTM (Finkbeiner et al., 1997; Kida, 2012; Silva et al., 1998; Suzuki et al., 2011). That is, by increasing the expression levels of various neurotrophins, CREB regulates memory formation, consolidation, and reconsolidation in a wide range of animal models, from invertebrates to mammals (Alberini, 2009, 2011; Bozon et al., 2003; Stevens, 1994; Yin and Tully, 1996). Therefore, it is not surprising that CREB and the signalling pathways leading to its activation are attractive targets for treatment aimed at improving memory function, in both diseased and healthy individuals (Barco et al., 2003; Khan et al., 2018; Tully et al., 2003).

Importantly for us here in the pond snail *Lymnaea stagnalis* (Linnaeus 1758) it was found that treatment with Q resulted in the upregulation of the orthologous gene of CREB1 (*Lym*CREB1) 3h after exposure to Q (Batabyal et al., 2021). In recent years, several studies from our lab have focused on how environmentally relevant stressors and bio-active compounds alter LTM formation in snails. Using *Lymnaea* as a model organism, it was shown for the first time in invertebrates that the flavonoid (-)- Epicatechin present in cocoa and green tea enhances LTM following operant conditioning of aerial respiratory behaviour (Fernell et al., 2016; Fruson et al., 2012; Knezevic and Lukowiak, 2014). Using operant conditioning of aerial respiratory behaviour it has been shown that memory formation can be altered (i.e., enhanced or inhibited) by a number of different factors (Lukowiak et al., 1996; Lukowiak et al., 2000, 2010). Consistent with the studies in vertebrate models (Debiec et al., 2002; Kida et al., 2002; Milekic and Alberini, 2002), new RNA and protein synthesis are required for LTM formation and reconsolidation in *Lymnaea* (Sangha, Scheibenstock, and Lukowiak, 2003).

Here we investigated the role of a plant flavonoid, Q, on memory formation, consolidation, and reconsolidation. Following configural learning, *Lymnaea* exposed to Q show an upregulation of *Lym*CREB1 which significatively enhanced LTM formation (Batabyal et al., 2021). Those results paved the way for our detailed study on the effects of Q on different LTM phases following operant conditioning of aerial respiratory behaviour. Here we assessed for the first time in an invertebrate model the effects of the exposure to Q on LTM formation, consolidation, and recall. We found that following the exposure of snails to Q 3h before or after training and 3h before a 24h memory test that LTM formation was enhanced.

## Material and methods

### Animals

*Lymnaea stagnalis* were bred and raised in the snail facility at the University of Calgary from a strain of *Lymnaea* originally obtained from Vrije Universiteit in Amsterdam. We have designated these snails as the W-strain of *Lymnaea* to identify them from the other strains used in our laboratory. Other strains of *Lymnaea* have different memory abilities following operant conditioning of aerial respiration (Sunada et al., 2017). Adult snails (with a shell length of 2.5–3.0 cm) were maintained at a density of one snail per litre in artificial pond water (PW) under eumoxic (i.e., normal O_2_ levels; PO_2_>9975 Pa) conditions. Artificial PW consisted of distilled water containing 26g l^−1^ Instant Ocean (Spectrum Brands, Madison, WI, USA), with added calcium sulphate dihydrate to create what we refer to as a ‘standard calcium level’ of 80 mg l^−1^ (Dalesman and Rundle, 2010; Knezevic et al., 2011). Animals were maintained at room temperature (20–22 °C) on a schedule of 16 h:8 h light: dark and had *ad libitum* access to romaine lettuce.

### Quercetin solution

Quercetin (3,3′,4′,5,7-pentahydroxyflavone, Q) was obtained from Sigma Chemical Company (St Louis, MO, USA) and was dissolved in 0.1% (final concentration) dimethyl sulfoxide (DMSO). We prepared a Q solution by dissolving 50μl in 500ml of pond water (PW).

### Aerial respiratory behaviour

*Lymnaea stagnalis* is a bimodal breather. In normal oxygenated conditions cutaneous respiration prevails; however, when in hypoxic conditions it switches to aerial respiration via the respiratory orifice, the pneumostome (Lukowiak et al., 1996; Lukowiak et al., 2006). Aerial respiration can be easily monitored via observation of the pneumostome opening and closing. Individually labeled snails were placed in 1 litre beaker filled with 500 ml of PW made hypoxic by vigorously bubbling N2 for 20 min (≤0.1 ml O_2_ l^−1^). Snails experiencing hypoxia increase their rate of aerial respiration (Lukowiak et al., 1996). Animals were acclimated for 10 min prior to observation. Bubbling was then reduced and continued at a slower rate during the following 30 min of breathing observation. In that way the environment was maintained hypoxic without disturbing snail activity. During the observation period aerial respiratory behaviour was recorded. In particular, we measured the amount of time the pneumostome was open for each snail without any tactile stimulation. This allowed us to compute the total breathing time in seconds for each snail. Following each breathing observation period, snails were returned to their home eumoxic aquaria.

### A 0.5h training session procedure for operant conditioning

In all of the operant conditioning training sessions and tests for LTM, a gentle tactile stimulus (using a sharpened wooden applicator) was applied to the pneumostome area (i.e., the respiratory orifice) every time the snail began to open its pneumostome to perform aerial respiration (Lukowiak et al., 1996). We refer to this response as an ‘attempted pneumostome opening’. This tactile stimulus only evoked pneumostome closure, without causing the animal to withdraw its foot and mantle area (i.e., the whole-animal withdrawal response) into its shell. Animals were given a 10 min acclimation period before the start of experiment, during which they could perform aerial respiration without receiving the negative reinforcement stimulus. The number of attempted pneumostome openings was recorded for each individual snail over the following 0.5h training session. To determine whether LTM was formed, an identical procedure (i.e., a memory test) was performed at different times following the training session. The time of the training session was defined as ‘T0’, whereas the memory tests were defined as times after the T0. Between the conditioning (training and memory) phases snails were returned to their eumoxic home aquaria, where they were fed *ad libitum* and freely allowed to perform aerial respiration. Breathing behaviour was not monitored while snails were in the home aquaria.

### Single poke procedure

Snails were placed in the hypoxic environment for a 10-min acclimation period without receiving any negative reinforcement stimulus. During the following 0.5h of exposure to hypoxic PW they received a poke only the first time they began to open the pneumostome. Hence the term ‘single poke procedure’ (Karnik et al., 2012).

### Statistical analysis

All our experimental treatments are independent from each other and have specific stimulus exposure timelines. Thus, we cannot combine the data in a meaningful manner for any grouped statistical analyses. We thus performed an individual paired-sample t test for each experiment. In all analysis reported here, a type I error rate of 0.05 was used. All statistical analyses were performed using GraphPad Prism v. 9.00e for MAC® (GraphPad Software, Inc., La Jolla, CA, USA).

## Results

### Effect of Q-exposure 3h before or after a 0.5h training session on LTM formation

We first determined if aerial respiratory behaviour was significantly influenced by Q. Thus, total breathing time (TBT) was measured in standard hypoxia in a cohort of snails (N = 10) 20h before and 3h after the exposure to Q for 1h. We found that TBT before and following exposure to Q was not significatively different (t = 0.72, df = 9, p = 0.49) (**Figure 1A**).

**Fig. 1.**
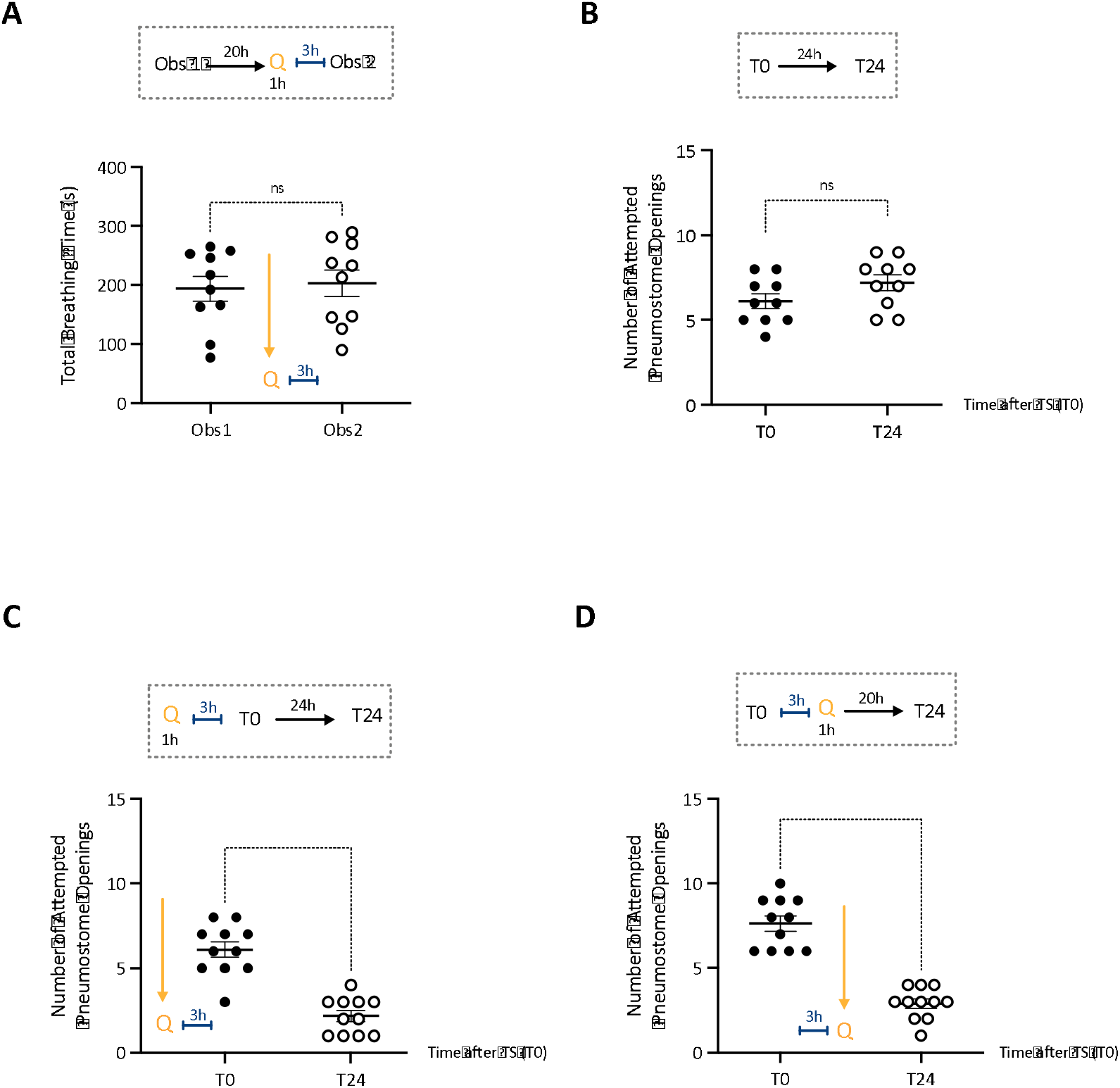
Quercetin exposure and aerial respiration. **(A)** Aerial respiratory behaviour in *Lymnaea* is not significantly altered by Q. We plotted and compared the total breathing time (TBT) between the first observation period (Obs1 – closed circles) and the second one performed 24h later, 3h after the exposure to Q (Obs2 – open circles). **(B)** Snails given a 0.5h TS in hypoxic PW do not form LTM. A naïve cohort of snails (N = 10) received a 0.5h training session (T0 – closed circles) and LTM was tested 24h later (T24 – open circles). **(C)** The exposure to Q 3h before a 0.5h training session enhances LTM formation. A naïve cohort of snails (N = 10) was exposed to Q for 1h and 3h later received a 0.5h training session (T0 – closed circles). 24h later animals were tested for LTM (T24 – open circles). **(D)** The exposure to Q 3h after a 0.5h training session enhances LTM formation. A naïve cohort of snails (N = 11) received a 0.5h training session (T0 – closed circles) and 3h after was exposed to Q for 1h. Snails were tested for LTM 24h later (T24 – open circles). The timeline for each experiment is presented above the data. The solid line is the mean, and the error bars are the s.e.m. Comparisons were made by paired t-test. ns = not significant; ****p<0.0001.

We next confirmed that in W-strain snails a 0.5h training session does not result in LTM formation (**Figure 1B**). We trained a naïve cohort of snails (N = 10) with a 0.5h training session (T0) and we tested for LTM 24h later (T24). These snails did not exhibit LTM as the number of attempted pneumostome openings at T0 and T24 was not statistically different (t = 1.94, df = 9; p = 0.084).

Then, we wished to determine if the Q-exposure 3h before training altered the ability of snails to form LTM (**Figure 1C**). Thus, a cohort of naïve snails (N = 11) was exposed to Q for 1h and three hours later received a 0.5h training session (T0). We tested memory 24h later (T24). Statistical analyses revealed that the number of attempted pneumostome openings at T24 was significantly reduced compared to T0 (t = 7.9; df = 10; p < 0.0001). Thus, Q-exposure 3h before the 0.5h training session enhanced LTM formation (i.e., LTM was present 24h after T0).

Finally, we asked whether Q-exposure 3h after a 0.5h training session (T0) would also enhance LTM formation (**Figure 1D**). A cohort of naïve snails (N = 11) was first trained in hypoxic PW (T0) and three hours later was placed for 1h in eumoxic Q-containing PW. Animals were then returned to their home aquaria and tested for LTM 20h later (i.e., at T24). Again, we found that at T24 the snails’ respiratory behaviour was significantly reduced with respect to T0 (t = 8.74; df = 10; p < 0.0001). Thus, a LTM lasting 24h was formed as a result of the exposure to Q after training.

Finally, we did control experiments to test the effect of the vehicle used to dissolve Q (i.e., DMSO) on LTM enhancement. We found that the exposure to DMSO 3h before the 0.5h training session did not result in LTM (T0: Mean ± sem = 6.90 ± 0.62; T24: Mean ± sem = 6.60 ± 0.60; t = 0.81, df = 9; p = 0.43). Similar results were obtained when DMSO exposure occurred 3h after the 0.5h training session (T0: Mean ± sem = 6.40 ± 0.54; T24: Mean ± sem = 6.0 ± 0.49; t = 0.82, df =9; p = 0.43).

Together these data show that Q-exposure enhanced LTM formation in W-strain snails when snails experience Q either 3h before or after a 0.5h training session. Exposing snails to vehicle only 3h before or 3h after training did not result in LTM. Finally, Q-exposure did not alter normal homeostatic aerial respiration. Thus, we concluded that Q exposure is capable of enhancing LTM formation.

An obvious experiment to perform was to train a naïve cohort of snails in hypoxic PW + Q and determine if this also resulted in enhanced LTM formation. Quite unexpectedly, we found that the combination of hypoxic PW and Q severely suppressed snails’ aerial respiratory behaviour, resulting in a sleep-like quiescent state that persisted for at least 2h after ending the exposure (data not shown). This phenomenon is currently under investigation in our lab.

### Effect of the single-poke procedure and Q-exposure on LTM

The demonstration that Q-exposure both 3h before and after a 0.5h training session procedure enhanced LTM formation and consolidation led us to investigate the effects of Q exposure on memory reconsolidation. Previous studies demonstrated that 24h after a training procedure that only results in an intermediate-term memory (ITM) a residual memory trace is present despite the LTM phenotype being absent (Parvez et al., 2005, 2006). Here (**Figure 2A**, N = 11), we found that a 0.5h training session followed by the ‘single poke’ procedure 24h after (i.e., T24) was not sufficient to enhance LTM at T48 (i.e., 48h after T0). In fact, there was no statistical difference between the number of attempted openings at T48 compared to T0 (t = 1.79, df = 10; p = 0.1).

**Fig. 2.**
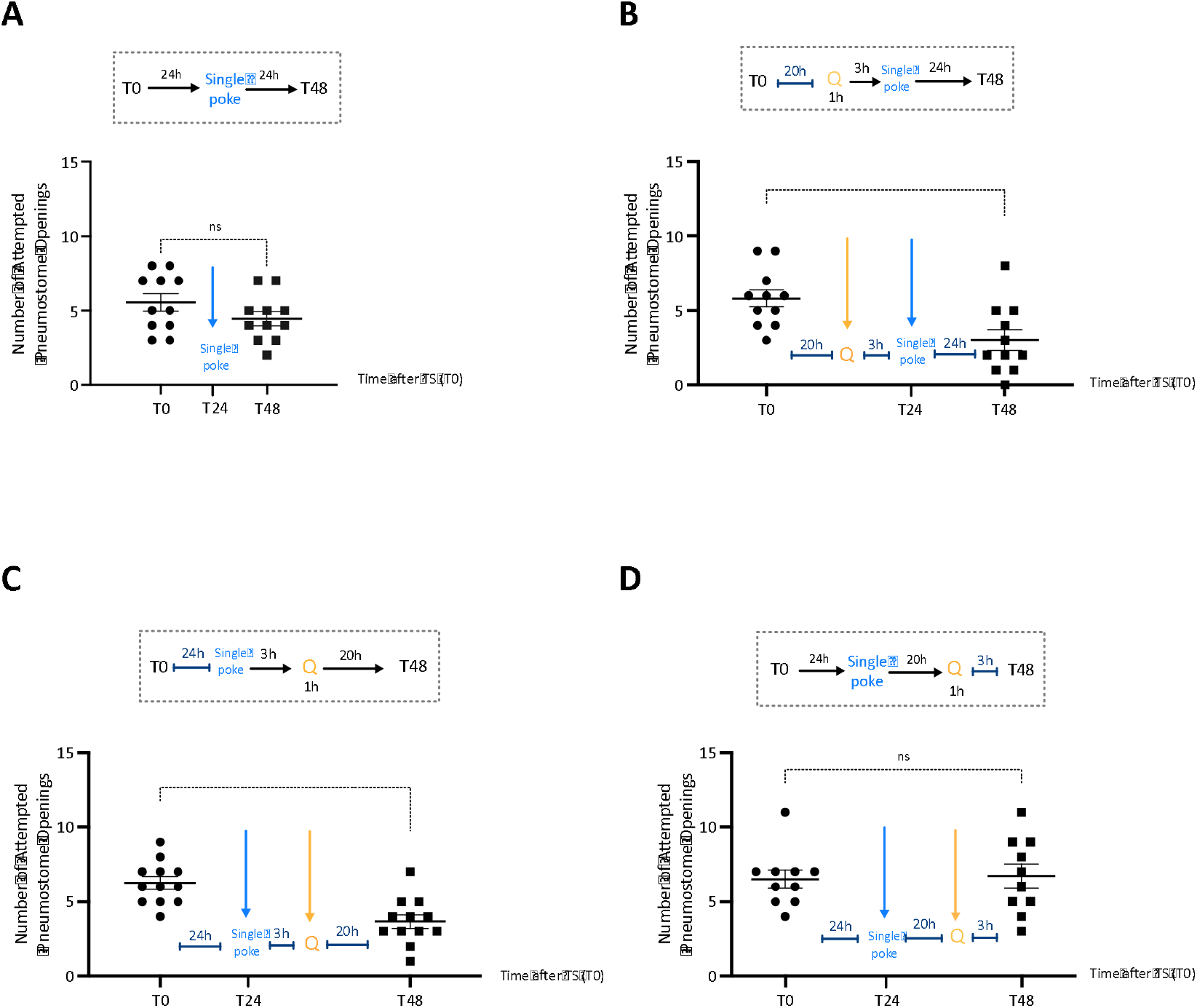
The single poke procedure, quercetin exposure, and enhanced LTM formation. **(A)** Snails receiving the single poke procedure 24h before the 0.5h training session do not form LTM. A naïve cohort of snails (N = 11) received a 0.5h training session (T0 – closed circles). 24h later animals received the ‘singe poke’ procedure and 24h later were tested for LTM (T48 – closed squares). **(B)** The exposure to Q 3h before the ‘single poke’ procedure occurring 24h after the 0.5h training session enhanced LTM formation. A naïve cohort of snails (N = 11) received a 0.5h training (T0 – closed circles) and 20h later was exposed to Q for 1h. 3h later snails received the single poke procedure and at T48 were tested for LTM (closed squares). **(C)** The exposure to Q 3h after the ‘single poke’ procedure occurring 24h after a 0.5h training session enhanced LTM formation. A naïve cohort of snails (N = 11) received a 0.5h training (T0 – closed circles) and 24h later was subjected to the single poke procedure. 3h later snails were exposed to Q for 1h and 20h later (at T48 – closed squares) were tested for LTM. **(D)** The exposure to Q 20h after the single poke procedure does not result in enhanced LTM formation. 24h after the 0.5h training session (T0 – closed circles), naïve snails (N = 10) were subjected to the single poke procedure. Q exposure occurred 20h later, 3h before LTM was tested at T48 (closed squares). The timeline for each experiment is presented above the data. The solid line is the mean, and the error bars are the s.e.m. Comparisons were made by paired t-test. ns = not significant; *p < 0.05; **** p < 0.0001.

We next asked (**Figure 2B**) whether the combined exposure to Q and the single poke procedure performed 24h after a 0.5h training session procedure (i.e., at T24) would result in LTM when memory was tested 24h later (i.e., T48). Thus, a naïve cohort of snails (N = 11) was first trained in a 0.5h training session (T0) and 20h later was exposed to Q for 1h. Three hours later (i.e., 24h after T0 – T24), animals were placed in the hypoxic environment and received a poke only the first time they began to open the pneumostome (i.e., ‘single poke’ procedure).

LTM was then tested 24h later (i.e., T48). In these snails a 48h LTM was shown (t = 3.02, df = 10; p = 0.012), since the number of attempted pneumostome openings at T48 was significantly less than that in T0. Thus, combining Q-exposure 3h before the ‘single-poke procedure’ resulted in an LTM when tested 48h after T0.

This result led us to hypothesize that Q-exposure 3h after the ‘single-poke’ procedure would also result in enhanced LTM formation. Thus, a naïve cohort of animals (N = 12) was tested using the same above behavioural procedure, but in these animals the exposure to Q occurred 3h after the ‘single poke’ procedure performed at T24 (**Figure 2C**). We found that at T48 the snails’ respiratory behaviour was significantly reduced with respect to T0 (t = 8.25, df = 11; p < 0.01), indicating that LTM persisted for at least 48h.

Finally, we showed in a cohort of snails (N = 10) trained as above that when Q-exposure was delayed by 20h after the ‘single-poke’ procedure, LTM was not present (**Figure 2D**). That is, the number of attempted pneumostome openings between the T0 and T48 was not significantly different (t = 0.3, df = 9; p=0.77). These data show that the ‘single poke’ procedure acts on a residual trace present 24h after T0 and when combined with Q-exposure 3h before or after, enhanced LTM formation.

### Effect of the single-poke procedure and Q-exposure before a 0.5h training session and LTM formation

Continuing with the effects of the ‘single poke’ procedure, we asked what effect, if any, the ‘single poke’ would have if the snails received it before a 0.5h training session. In a naïve cohort of snails (**Figure 3A**; N = 10) the single poke procedure 24h before T0 did not result in LTM when tested 24h later (i.e., T24). A paired t-test revealed that the number of attempted pneumostome openings at T24 was not significatively different from those at T0 (t = 0.58, df = 9, p = 0.59). Thus, LTM was not present. We then asked whether Q exposure 24h before a 0.5h training session (i.e., T0) resulted in enhanced LTM formation. Thus, a naïve cohort of snails (N = 10) was exposed to Q 24h before T0 and was then tested for LTM 24h later (**Figure 3B**). A paired t-test revealed that the number of attempted pneumostome openings at T24 was not significatively different from those at T0 (t = 1.33, df = 9, p = 0.21). That is, Q-exposure 24h before a 0.5h training session did not result in LTM formation.

**Fig. 3.**
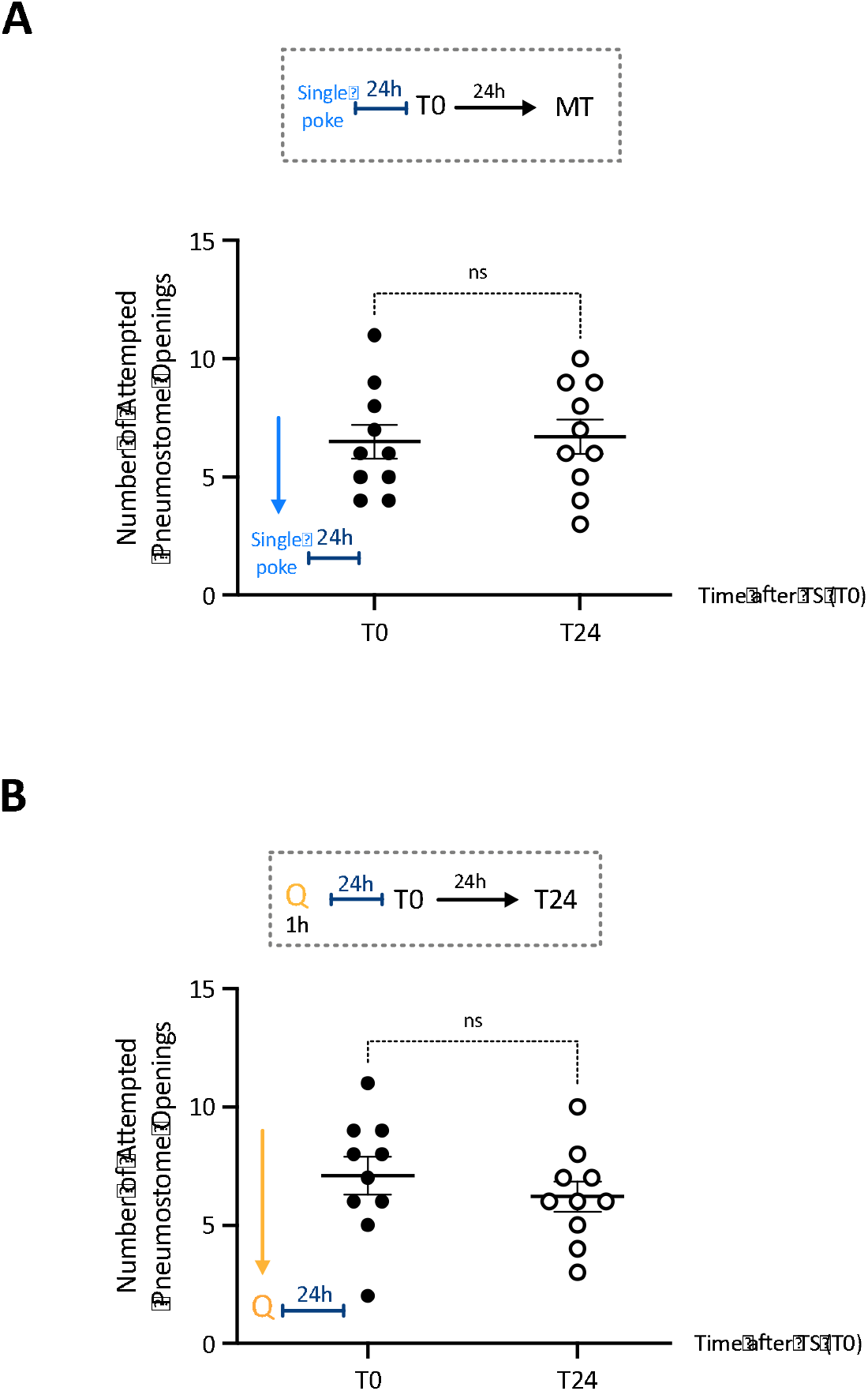
The single poke procedure or Q-exposure 24h before a 0.5h training session does not result in LTM formation. **(A)** When applied 24h before a 0.5h training session, the single poke procedure does not result in LTM formation. Snails were trained (T0 – closed circles) 24h after receiving the single-poke procedure and LTM was tested at T24 (open circles). **(B)** When applied 24h before the 0.5h training session, Q-exposure does not result in LTM formation. Snails were exposed to Q for 1h and 24h later received a 0.5h training session (T0 – closed circles). LTM was tested at T24 (open circles). The timeline of each experiment is presented above the data. Comparisons were made by paired t-test. The solid line is the mean, and the error bars are the s.e.m. ns = not significant.

We next asked if a combination of the ‘single poke’ procedure and Q-exposure 24h before a 0.5h training session (i.e., T0) would result in enhanced LTM formation. These data are presented in **Figure 4.** A naïve cohort of snails (N = 10) was first exposed to Q for 1h and 3h later was subjected to the ‘single poke’ procedure (**Figure 4A**). Animals were then returned to their home aquaria for 20h before being trained with a 0.5h training session (T0). These snails were then tested for LTM 24h later (i.e., T24). We found that the number of attempted openings at T24 was significantly less than at T0 (t = 7.23; df = 11; p < 0.0001). Thus, LTM was present. These data show that Q-exposure, combined with the single poke procedure 24h before training, resulted in LTM formation. We next presented to a naïve cohort of snails (N = 10) the single poke procedure first and then 3h later we exposed the snails to Q for 1h (**Figure 4B**). These snails then were trained with a 0.5h training session (T0) and were tested for LTM 24h later (i.e., T24). The sequential presentation of the single poke and then Q exposure resulted in LTM. That is, the number of attempted pneumostome openings in T24 was significantly less than in T0 (t = 2.84, df = 9; p = 0.02). Thus, the ‘single-poke’ procedure and Q-exposure 3h later was sufficient to cause LTM formation following a 0.5h training session 20h later. Not shown here, are control experiments where the vehicle is used rather than Q along with the single-poke procedure; in all cases enhanced LTM formation was not observed.

**Fig. 4.**
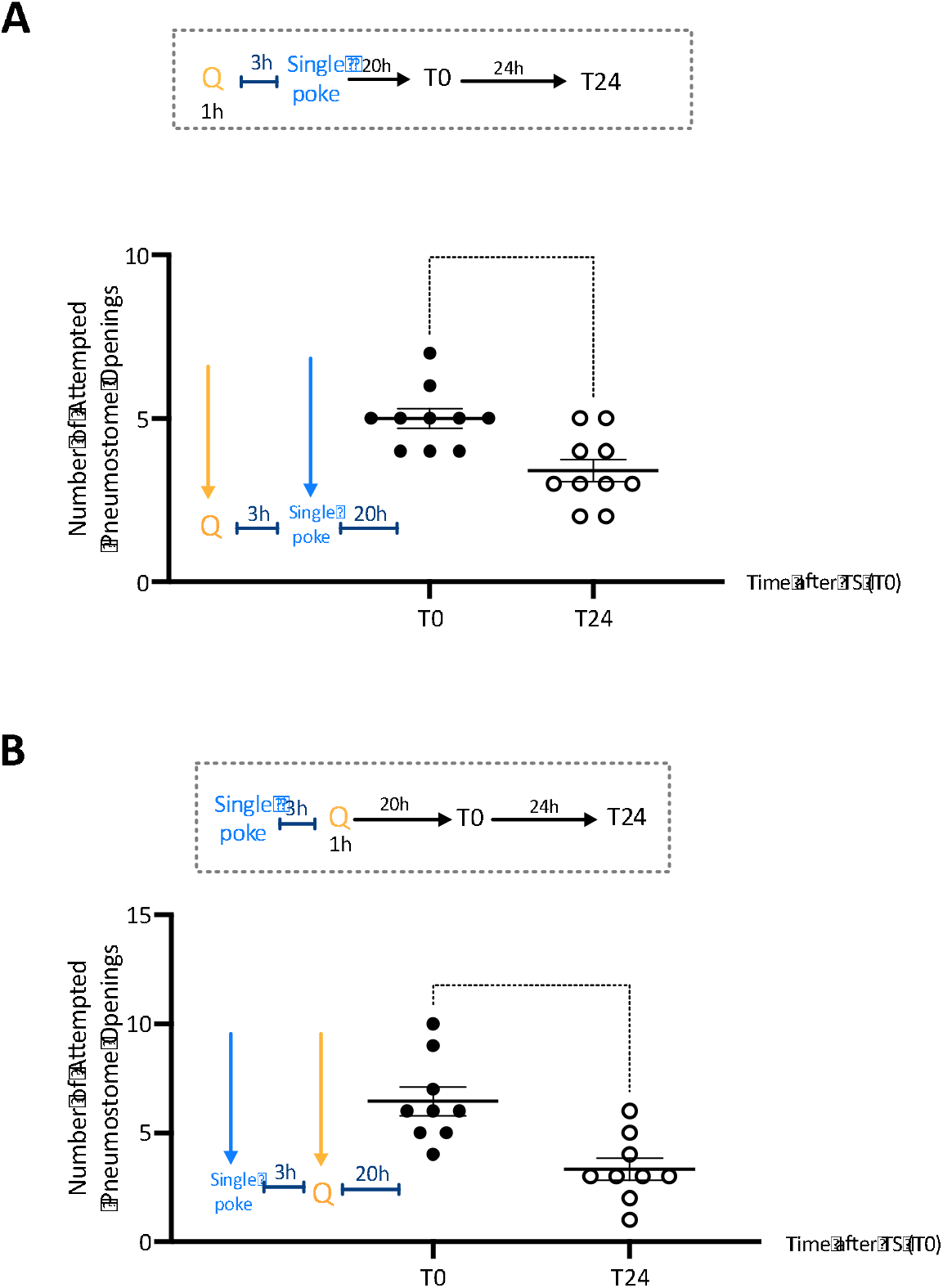
The single poke procedure combined with Q-exposure 3h before a 0.5h training session results in enhanced LTM formation. **(A)** The exposure to Q 3h before the single poke procedure performed before the single training results in LTM. A naïve cohort of snails (N = 10) was exposed to Q for 1h and 3h later received the single poke procedure. 20h later, animals were trained (T0 – closed circles) and LTM was tested at 24h (T24 – open circles). **(B)** The exposure to Q 3h after the single poke procedure performed before the single training results in LTM. A naïve cohort of snails (N = 9) received the single poke procedure and 3h later was exposed to Q for 1h. 20h later, animals were trained (T0 – closed circles) and LTM was tested at T24 (open circles). The timeline of each experiment is presented above the data. Comparisons were made by paired t-test. **** p < 0.0001; * p < 0.05.

## Discussion

This study tested the hypothesis that Q-exposure during critical periods of memory formation enhanced LTM formation following operant conditioning of aerial respiratory behaviour. Thus, Q-exposure 3h before or after the different phases of learning and memory (e.g., consolidation and recall) lead to enhance LTM formation. In addition, Q-exposure improved LTM formation following the ‘single poke’ procedure. These data are all consistent with Q-exposure upregulating *LymCREB1* which then synergistically interacts with the other molecular events necessary to form LTM. Q-exposure has previously been shown to act in the snails’ nervous system to both block the up-regulation of heat shock proteins (HSPs) following a heat shock and to up-regulate *LymCREB1* (Foster et al., 2015; Sunada et al., 2016; Rivi et al., 2021a; Batabyal et al., 2021). Since *LymCREB1* plays a key role in conditioned taste aversion memory formation in *Lymnaea* (Sadamoto et al., 2004), our data extend those findings to include memory formation following operant conditioning of aerial respiratory behaviour. Here we showed that Q-exposure alters four different phases of memory: 1) acquisition (i.e., a learning event), 2) consolidation processes after acquisition, 3) memory recall, and 4) memory reconsolidation. In all these memory phases Q-exposure enhanced LTM persistence.

Memory is stored in specific neural networks termed ‘memory engrams’ (Semon, 1921) that can be modified by factors such as stress, increased training and, in the data shown here, the exposure to certain bio-active products such as Q. While not examined here, we hypothesize that Q-exposure alters the activity of an identified neuron, RPeD1 (Right Pedal Dorsal 1), known to be a necessary site for LTM formation, extinction, and reconsolidation (Lukowiak et al., 2006; Sangha et al., 2005; Sangha et al., 2003a,b; Sangha, Scheibenstock, and Lukowiak, 2003; Scheibenstock et al., 2002). Previously, it has been shown that predator detection (using crayfish effluent – CE) alters the behaviour of this neuron leading to enhanced LTM formation (Orr and Lukowiak, 2008). However, for that to occur snails had to be trained in CE. The exposure to CE 1h before or 1h after a single 0.5h did not cause LTM enhancement. In a similar vein, another bio-active substance, epicatechin, also had to be experienced either during or immediately after training to cause LTM enhancement (Fernell et al., 2016)

Our working hypothesis was that Q-exposure, which causes an up-regulation of *LymCREB1* 3 hours after exposure (Batabyal et al., 2021), results in enhanced LTM formation when Q-exposure occurs within 3 hours of the different phases of learning and memory formation. That is, boosting *LymCREB1* activity during the 3-hour window prior to or after the single 0.5h operant conditioning training session may amplify the on-going and necessary molecular processes that underlie LTM. The experiments reported here were all designed to test this hypothesis. Our data also add to our knowledge that bio-active, naturally occurring substances, such as Q, can alter cognitive ability (e.g., enhanced LTM formation). Finally, the data also illustrate the advantages of using our model system in gaining an understanding of how substances, such as Q, alter cognition. Q, as well as many other compounds (e.g., epicatechin), easily penetrate through the integument of the snail and, because snails have an open circulatory system, can easily and quickly reach the CNS (Fruson et al., 2012; Rivi et al., 2021b, Rivi et al., 2020).

The highly conserved CREB gene and CREB-like proteins have been characterized in *Lymnaea* nervous system (Frank and Greenberg, 1994) Ribeiro et al., 2003; Sadamoto et al., 2004) and it has been demonstrated that the conditioned taste aversion (CTA) training increased *LymCREB1* expression levels (Sadamoto et al., 2010). Moreover, the levels of phosphorylated CREB1 are also increased following appetitive classical conditioning of feeding behaviour (Ribeiro et al., 2003). In particular, in appetitive conditioning studies it was demonstrated that between 10 min and 1 h after training, many translational-dependent processes are activated to form ITM. While ITM only requires the synthesis of new proteins from pre-existing mRNA transcripts, LTM is dependent on altered gene activity, new protein synthesis and transport of the protein(s) to the site of plasticity (Braun and Lukowiak, 2011; Lukowiak et al., 2000; Rosenzweig et al., 1993). ITM and LTM are independent but interrelated events which occur in parallel and has specific temporal phases and transition periods (Marra et al., 2013). As CREB1 is a regulator of mRNA transcription, it most likely does not play a necessary role in ITM formation but does so for LTM formation (Sadamoto et al., 2004). Studies in *Drosophila* and rodent models also demonstrate that neurons with higher CREB levels are preferentially recruited for LTM engrams as *de novo* transcription and translation is required during formation and consolidation of LTM (Han et al., 2007; Miyashita et al., 2018).

In the W-strain of *Lymnaea* used here a 0.5h training session is only capable of causing a memory that persists for approximately 3h and this has been termed intermediate-term memory (ITM, Sangha et al., 2003a,b). However, Parvez et al., (2005, 2006) found that there was a residual memory trace 24h after training, even though the LTM phenotype was absent. Thus, it was apparent to us that we could ascertain whether the exposure to Q 3h before or after this period caused LTM enhancement when combined with the presentation of a stimulus associated with learning (i.e., the single poke procedure). As we showed here, exposing snails to Q 20h after a 0.5h training session was sufficient for the single-poke procedure performed 3h later to cause LTM formation when tested after T0. Similarly, we showed that using the single-poke procedure 24h after the initial 0.5h training session and then 3h later exposing snails to Q resulted in a LTM when tested 48h after the initial training session (i.e., T0). Both the consolidation and reconsolidation processes of LTM in our model system, as well as in vertebrates, have been shown to be dependent on altered gene activity (i.e., transcription) and *de novo* protein synthesis (i.e., translation) (Anokhin et al., 2002; Debiec et al., 2002; Judge and Quartermain, 1982; Kida et al., 2002; Milekic and Alberini, 2002; Nader and Einarsson, 2010). Moreover, there are several lines of evidence implying that memory consolidation and reconsolidation use similar molecular pathways that converge on the activation of CREB1 (Alberini, 2011; McKenzie and Eichenbaum, 2011; Nader and Einarsson, 2010; Tronson and Taylor, 2007).

Together these data are consistent with our hypothesis that Q, which induces an upregulation of *LymCREB1*, interacts synergistically with the molecular processes that underlie LTM formation, consolidation, and reconsolidation. Thus, Q-exposure, in essence, boosts those processes if the boost occurs within a specific ‘window of opportunity’. Outside of that window, an enhancement of LTM formation is not observed.

We further showed, using the single-poke procedure, that the exposure to Q 3h before or after the single poke procedure enabled a 0.5h training session, given some 20h after the Q-exposure combined with the single-poke procedure, to result in LTM formation. These data suggest that the Q-induced upregulation of *LymCREB1* persists for sufficient time, such that the one 0.5h training session became capable of inducing the molecular changes necessary for LTM formation. We plan to investigate this in detail in our future experiments.

Our findings reported here on the enhancement of LTM formation with Q-exposure are also consistent with our recent findings on configural learning in *Lymnaea* (Batabyal et al., 2021). In that study, Q-exposure 3h prior to, immediately before or 3h following configural learning training, enhanced LTM formation. Usually, the configural learning procedure only results in memory persisting for 3h (i.e., ITM) but following Q-exposure an LTM was formed that persisted in some cases for 48h (Batabyal et al., 2021).

Previous studies demonstrated that a 0.5h is sufficient to produce an ITM trace lasting 3h, which is dependent on new protein synthesis (Braun and Lukowiak, 2011; Lukowiak et al., 2000; Sangha et al., 2003a,b). It is at this 3h post training time, in appetitive food conditioning, that *Lymnaea* show a transition from ‘late-ITM’ to LTM that requires RNA synthesis (Marra et al., 2013). During the 3h interval between training and Q-exposure the molecular machinery required for ITM to be converted to LTM is induced by Q via upregulation of *LymCREB1*. These data are consistent with previous studies from both invertebrates and vertebrates showing the key role of CREB1 in LTM formation and consolidation after various learning paradigms (Alberini, 2011; Sangha, Scheibenstock, and Lukowiak, 2003). We conclude that Q-exposure 3h before or after training, while acting on different mnemonic phases (i.e., memory formation and consolidation, respectively), enhances the persistence of LTM in *Lymnaea*.

Our data are also consistent with the hypothesis that dietary sources of Q could improve cognitive ability (Finkbeiner et al., 1997; Kida, 2012; Silva et al., 1998; Suzuki et al., 2011). Further, our data illustrate that *Lymnaea* is a suitable model to elucidate neuronal and molecular mechanisms induced by Q-exposure during the different phases in the process of transforming learning into memory and its recall. We hypothesize that *LymCREB1* activation is differentially involved and regulated depending on the time of Q-exposure. Other studies suggested that flavonoids may modulate kinase activity [e.g., mitogen-activated protein kinase (MAPK) cascade] and signalling cascades lying downstream of these kinases (Spencer, 2007). The protein kinase/MAPK signalling cascades have been shown to be necessary for both ITM and LTM formation in *Lymnaea* (Ribeiro et al., 2003). Therefore, alteration of protein kinase activity following Q-exposure may potentially be the mechanism by which memory formation is enhanced in *Lymnaea*.

Findings presented here provide the groundwork for future molecular analysis of how Q-exposure acts at the neuronal level and the mechanisms involved in the alterations to memory formation, storage and, recall, paving the way for interesting translational neuroscience studies. Our current data provides the first support for Q-modulated enhancement of cognitive function in an invertebrate model after an operant conditioning procedure. Thus, strengthening the case for *Lymnaea* as a model for translational neuroscience (Rivi et al., 2021b, Fodor et al., 2020).

## Acknowledgements

We would like to thank Diana Kagan, Bevin Wiley, and David Chau for the discussions during the research.

## Funding

Funding was provided by the Natural Sciences and Engineering Research Council of Canada (NSERC) to KL.

## Competing interests

We have no competing interests.

## Author’s contributions

VR, AB, and KL conceptualised and designed study. VR performed behavioural data collection, analysis and writing first draft. AB, CB, JMCB, and FT critically revised manuscript. KL coordinated study, provided funding, and critically revised manuscript. All authors take responsibility of the data provided and consent to the publication of the manuscript.

## Notes

### Competing Interest Statement

The authors have declared no competing interest.

## References

Ahmad, S., Ullah, F., Ayaz, M., Sadiq, A. and Imran, M. (2015). Antioxidant and anticholinesterase investigations of Rumex hastatus D. Don: Potential effectiveness in oxidative stress and neurological disorders. Biological Research, 48(1), 20. https://doi.org/10.1186/s40659-015-0010-2

Alberini, C. M. (2009). Transcription Factors in Long-Term Memory and Synaptic Plasticity. Physiological Reviews, 89(1). https://doi.org/10.1152/physrev.00017.2008

Alberini, C. M. (2011). The Role of Reconsolidation and the Dynamic Process of Long-Term Memory Formation and Storage. Frontiers in Behavioural Neuroscience, 5. https://doi.org/10.3389/fnbeh.2011.00012

Anokhin, K. V., Tiunova, A. A. and Rose, S. P. R. (2002). Reminder effects – reconsolidation or retrieval deficit? Pharmacological dissection with protein synthesis inhibitors following reminder for a passive-avoidance task in young chicks: Memory consolidation and reminder effects in chicks. European Journal of Neuroscience, 15(11), 1759–1765. https://doi.org/10.1046/j.1460-9568.2002.02023.x

Ansari, M. A., Abdul, H. M., Joshi, G., Opii, W. O. and Butterfield, D. A. (2009). Protective effect of quercetin in primary neurons against Aβ(1–42): Relevance to Alzheimer’s disease. The Journal of Nutritional Biochemistry, 20(4), 269–275. https://doi.org/10.1016/j.jnutbio.2008.03.002

Ayaz, M., Sadiq, A., Junaid, M., Ullah, F., Ovais, M., Ullah, I., Ahmed, J. and Shahid, M. (2019). Flavonoids as Prospective Neuroprotectants and Their Therapeutic Propensity in Aging Associated Neurological Disorders. Frontiers in Aging Neuroscience, 11. https://doi.org/10.3389/fnagi.2019.00155

Bakoyiannis, I., Daskalopoulou, A., Pergialiotis, V. and Perrea, D. (2019). Phytochemicals and cognitive health: Are flavonoids doing the trick? Biomedicine and Pharmacotherapy, 109, 1488–1497. https://doi.org/10.1016/j.biopha.2018.10.086

Barco, A., Pittenger, C. and Kandel, E. R. (2003). CREB, memory enhancement and the treatment of memory disorders: Promises, pitfalls and prospects. Expert Opinion on Therapeutic Targets, 7(1), 101–114. https://doi.org/10.1517/14728222.7.1.101

Batabyal, A., Rivi, V., Benatti, C., Blom, J. M. C., and Lukowiak, K. (2021). Long term memory of configural learning is enhanced via CREB upregulation by the flavonoid Quercetin in *Lymnaea stagnalis*. Journal of Experimental Biology. J Exp Biol jeb.242761. https://doi.org/10.1242/jeb.242761

Bhutada, P., Mundhada, Y., Bansod, K., Bhutada, C., Tawari, S., Dixit, P., and Mundhada, D. (2010). Ameliorative effect of quercetin on memory dysfunction in streptozotocin-induced diabetic rats. Neurobiology of Learning and Memory, 94(3), 293–302. https://doi.org/10.1016/j.nlm.2010.06.008

Bisaz, R., Travaglia, A., and Alberini, C. M. (2014). The neurobiological bases of memory formation: From physiological conditions to psychopathology. Psychopathology, 47(6), 347–356. https://doi.org/10.1159/000363702

Bozon, B., Kelly, Á., Josselyn, S. A., Silva, A. J., Davis, S. and Laroche, S. (2003). MAPK, CREB and *zif268* are all required for the consolidation of recognition memory. Philosophical Transactions of the Royal Society of London. Series B: Biological Sciences, 358(1432), 805–814. https://doi.org/10.1098/rstb.2002.1224

Braun, M. H. and Lukowiak, K. (2011). Intermediate and long-term memory are different at the neuronal level in Lymnaea stagnalis (L.). Neurobiology of Learning and Memory, 96(2), 403–416. https://doi.org/10.1016/j.nlm.2011.06.016

Brem, A.-K., Ran, K. and Pascual-Leone, A. (2013). Learning and memory. Handbook of Clinical Neurology, 116, 693–737. https://doi.org/10.1016/B978-0-444-53497-2.00055-3

Broman-Fulks, J. J., Canu, W. H., Trout, K. L. and Nieman, D. C. (2012). The effects of quercetin supplementation on cognitive functioning in a community sample: A randomized, placebo-controlled trial. Therapeutic Advances in Psychopharmacology, 2(4), 131–138. https://doi.org/10.1177/2045125312445894

Conte, C., Herdegen, S., Kamal, S., Patel, J., Patel, U., Perez, L., Rivota, M., Calin-Jageman, R. J. and Calin-Jageman, I. E. (2017). Transcriptional correlates of memory maintenance following long-term sensitization of Aplysia californica. Learning and Memory (Cold Spring Harbor, N.Y.), 24(10), 502–515. https://doi.org/10.1101/lm.045450.117

Cushnie, T. P. T. and Lamb, A. J. (2005). Antimicrobial activity of flavonoids. International Journal of Antimicrobial Agents, 26(5), 343–356. https://doi.org/10.1016/j.ijantimicag.2005.09.002

Dalesman, S. and Rundle, S. (2010). Influence of rearing and experimental temperatures on predator avoidance behaviour in a freshwater pulmonate snail. Freshwater Biology, 55, 2107–2113. https://doi.org/10.1111/j.1365-2427.2010.02470.x

Davis, R. L. (2011). Traces of Drosophila Memory. Neuron, 70(1), 8–19. https://doi.org/10.1016/j.neuron.2011.03.012

Debiec, J., LeDoux, J. E. and Nader, K. (2002). Cellular and Systems Reconsolidation in the Hippocampus. Neuron, 36(3), 527–538. https://doi.org/10.1016/S0896-6273(02)01001-2

Eggler, A. L., Gay, K. A. and Mesecar, A. D. (2008). Molecular mechanisms of natural products in chemoprevention: Induction of cytoprotective enzymes by Nrf2. Molecular Nutrition and Food Research. https://doi.org/10.1002/mnfr.200700249

Fernell, M., Swinton, C., and Lukowiak, K. (2016). Epicatechin, a component of dark chocolate, enhances memory formation if applied during the memory consolidation period. Communicative and Integrative Biology, 9(4), e1205772. https://doi.org/10.1080/19420889.2016.1205772

Finkbeiner, S., Tavazoie, S. F., Maloratsky, A., Jacobs, K. M., Harris, K. M. and Greenberg, M. E. (1997). CREB: A Major Mediator of Neuronal Neurotrophin Responses. Neuron, 19(5), 1031–1047. https://doi.org/10.1016/S0896-6273(00)80395-5

Fodor, I., Hussein, A. A., Benjamin, P. R., Koene, J. M., and Pirger, Z. (2020). The unlimited potential of the great pond snail, Lymnaea stagnalis. ELife, 9, e56962. https://doi.org/10.7554/eLife.56962

Foster, N. L., Lukowiak, K. and Henry, T. B. (2015). Time-related expression profiles for heat shock protein gene transcripts (HSP40, HSP70) in the central nervous system of Lymnaea stagnalis exposed to thermal stress. Communicagive and integrative biology, 8(3), e1040954. https://doi.org/10.1080/19420889.2015.1040954

Frank, D. A. and Greenberg, M. E. (1994). CREB: A mediator of long-term memory from mollusks to mammals. Cell, 79(1), 5–8. https://doi.org/10.1016/0092-8674(94)90394-8

Fruson, L., Dalesman, S. and Lukowiak, K. (2012). A flavonol present in cocoa [(-)epicatechin] enhances snail memory. Journal of Experimental Biology, 215(20), 3566–3576. https://doi.org/10.1242/jeb.070300

Haleagrahara, N., Siew, C. J., Mitra, N. K. and Kumari, M. (2011). Neuroprotective effect of bioflavonoid quercetin in 6-hydroxydopamine-induced oxidative stress biomarkers in the rat striatum. Neuroscience Letters, 500(2), 139–143. https://doi.org/10.1016/j.neulet.2011.06.021

Han, J.-H., Kushner, S. A., Yiu, A. P., Cole, C. J., Matynia, A., Brown, R. A., Neve, R. L., Guzowski, J. F., Silva, A. J. and Josselyn, S. A. (2007). Neuronal Competition and Selection During Memory Formation. Science, 316(5823), 457–460. https://doi.org/10.1126/science.1139438

Havsteen, B. H. (2002). The biochemistry and medical significance of the flavonoids. Pharmacology and Therapeutics, 96(2–3), 67–202. https://doi.org/10.1016/s0163-7258(02)00298-x

Hawkins, R. D., Kandel, E. R. and Bailey, C. H. (2006). Molecular mechanisms of memory storage in Aplysia. The Biological Bulletin, 210(3), 174–191. https://doi.org/10.2307/4134556

Isabel, G., Pascual, A. and Preat, T. (2004). Exclusive consolidated memory phases in Drosophila. Science (New York, N.Y.), 304(5673), 1024–1027. https://doi.org/10.1126/science.1094932

Jayasena, T., Poljak, A., Smythe, G., Braidy, N., Münch, G. and Sachdev, P. (2013). The role of polyphenols in the modulation of sirtuins and other pathways involved in Alzheimer’s disease. Ageing Research Reviews, 12(4), 867–883. https://doi.org/10.1016/j.arr.2013.06.003

Judge, M. and Quartermain, D. (1982). Characteristics of retrograde amnesia following reactivation of memory in mice. Physiology and Behaviour, 28(4), 585–590. https://doi.org/10.1016/0031-9384(82)90034-8

Kandel, E. R., Dudai, Y. and Mayford, M. R. (2014). The molecular and systems biology of memory. Cell, 157(1), 163–186. https://doi.org/10.1016/j.cell.2014.03.001

Karimipour, M., Rahbarghazi, R., Tayefi, H., Shimia, M., Ghanadian, M., Mahmoudi, J. and Bagheri, H. S. (2019). Quercetin promotes learning and memory performance concomitantly with neural stem/progenitor cell proliferation and neurogenesis in the adult rat dentate gyrus. International Journal of Developmental Neuroscience: The Official Journal of the International Society for Developmental Neuroscience, 74, 18–26. https://doi.org/10.1016/j.ijdevneu.2019.02.005

Karnik, V., Dalesman, S. and Lukowiak, K. (2012). Input from a chemosensory organ, the osphradium, does not mediate aerial respiration in Lymnaea stagnalis. Aquatic Biology, 15(2), 167–173. https://doi.org/10.3354/ab00416

Khan, H., Marya, Amin, S., Kamal, M. A., and Patel, S. (2018). Flavonoids as acetylcholinesterase inhibitors: Current therapeutic standing and future prospects. Biomedicine and Pharmacotherapy, 101, 860–870. https://doi.org/10.1016/j.biopha.2018.03.007

Kida, S. (2012). A Functional Role for CREB as a Positive Regulator of Memory Formation and LTP. Experimental Neurobiology, 21(4), 136–140. https://doi.org/10.5607/en.2012.21.4.136

Kida, S., Josselyn, S. A., de Ortiz, S. P., Kogan, J. H., Chevere, I., Masushige, S., and Silva, A. J. (2002). CREB required for the stability of new and reactivated fear memories. Nature Neuroscience, 5(4), 348–355. https://doi.org/10.1038/nn819

Knezevic, B., Dalesman, S., Karnik, V., Byzitter, J., and Lukowiak, K. (2011). Low external environmental calcium levels prevent forgetting in Lymnaea. Journal of Experimental Biology, 214(12), 2118–2124. https://doi.org/10.1242/jeb.054635

Knezevic, B., and Lukowiak, K. (2014). The flavonol epicatechin reverses the suppressive effects of a stressor on long-term memory formation. Journal of Experimental Biology, 217(22), 4004–4009. https://doi.org/10.1242/jeb.110726

Kong, A.-N. T., Yu, R., Chen, C., Mandlekar, S., and Primiano, T. (2000). Signal transduction events elicited by natural products: Role of MAPK and caspase pathways in homeostatic response and induction of apoptosis. Archives of Pharmacal Research, 23(1), 1–16. https://doi.org/10.1007/BF02976458

Letenneur, L., Proust-Lima, C., Le Gouge, A., Dartigues, J., and Barberger-Gateau, P. (2007). Flavonoid Intake and Cognitive Decline over a 10-Year Period. American Journal of Epidemiology, 165(12), 1364–1371. https://doi.org/10.1093/aje/kwm036

Lukowiak, K., Ringseis, E., Spencer, G., Wildering, W., and Syed, N. (1996). Operant conditioning of aerial respiratory behaviour in Lymnaea stagnalis. The Journal of Experimental Biology, 199(Pt 3), 683–691.

Lukowiak, K., Adatia, N., Krygier, D. and Syed, N. (2000). Operant Conditioning in Lymnaea: Evidence for Intermediate- and Long-term Memory. Learning and Memory, 7(3), 140–150.

Lukowiak, K., Martens, K., Orr, M., Parvez, K., Rosenegger, D. and Sangha, S. (2006). Modulation of aerial respiratory behaviour in a pond snail. Respiratory Physiology and Neurobiology, 154(1–2), 61–72. https://doi.org/10.1016/j.resp.2006.02.009

Lukowiak, K., Orr, M., de Caigny, P., Lukowiak, K. S., Rosenegger, D., Han, J. I. and Dalesman, S. (2010). Ecologically relevant stressors modify long-term memory formation in a model system. Behavioural Brain Research, 214(1), 18–24. https://doi.org/10.1016/j.bbr.2010.05.011

Macready, A. L., Kennedy, O. B., Ellis, J. A., Williams, C. M., Spencer, J. P. E. and Butler, L. T. (2009). Flavonoids and cognitive function: A review of human randomized controlled trial studies and recommendations for future studies. Genes and Nutrition, 4(4), 227–242. https://doi.org/10.1007/s12263-009-0135-4

Manach, C., Scalbert, A., Morand, C., Rémésy, C. and Jiménez, L. (2004). Polyphenols: Food sources and bioavailability. The American Journal of Clinical Nutrition, 79(5), 727–747. https://doi.org/10.1093/ajcn/79.5.727

Manach, C., Williamson, G., Morand, C., Scalbert, A. and Rémésy, C. (2005). Bioavailability and bioefficacy of polyphenols in humans. I. Review of 97 bioavailability studies. The American Journal of Clinical Nutrition, 81(1), 230S–242S. https://doi.org/10.1093/ajcn/81.1.230S

Mandel, S. and Youdim, M. B. H. (2004). Catechin polyphenols: Neurodegeneration and neuroprotection in neurodegenerative diseases. Free Radical Biology and Medicine, 37(3), 304–317. https://doi.org/10.1016/j.freeradbiomed.2004.04.012

Marra, V., O’Shea, M., Benjamin, P. and Kemenes, I (2013) Susceptibility of memory consolidation during lapses in recall. Nat Comm 4:1578

Mastroiacovo, D., Kwik-Uribe, C., Grassi, D., Necozione, S., Raffaele, A., Pistacchio, L., Righetti, R., Bocale, R., Lechiara, M. C., Marini, C. et al., (2015). Cocoa flavanol consumption improves cognitive function, blood pressure control, and metabolic profile in elderly subjects: The Cocoa, Cognition, and Aging (CoCoA) Study—a randomized controlled trial. The American Journal of Clinical Nutrition, 101(3), 538–548. https://doi.org/10.3945/ajcn.114.092189

Matsuno, M., Horiuchi, J., Yuasa, Y., Ofusa, K., Miyashita, T., Masuda, T. and Saitoe, M. (2015). Long-Term Memory Formation in Drosophila Requires Training-Dependent Glial Transcription. The Journal of Neuroscience, 35(14), 5557–5565. https://doi.org/10.1523/JNEUROSCI.3865-14.2015

McKenzie, S., and Eichenbaum, H. (2011). Consolidation and reconsolidation: Two lives of memories? Neuron, 71(2), 224–233. https://doi.org/10.1016/j.neuron.2011.06.037

Milekic, M. H. and Alberini, C. M. (2002). Temporally Graded Requirement for Protein Synthesis following Memory Reactivation. Neuron, 36(3), 521–525. https://doi.org/10.1016/S0896-6273(02)00976-5

Milner, B., Squire, L. R. and Kandel, E. R. (1998). Cognitive neuroscience and the study of memory. Neuron, 20(3), 445–468. https://doi.org/10.1016/s0896-6273(00)80987-3

Miyashita, T., Kikuchi, E., Horiuchi, J. and Saitoe, M. (2018). Long-Term Memory Engram Cells Are Established by c-Fos/CREB Transcriptional Cycling. Cell Reports, 25(10), 2716–2728.e3. https://doi.org/10.1016/j.celrep.2018.11.022

Morel, I., Lescoat, G., Cogrel, P., Sergent, O., Pasdeloup, N., Brissot, P., Cillard, P., and Cillard, J. (1993). Antioxidant and iron-chelating activities of the flavonoids catechin, quercetin and diosmetin on iron-loaded rat hepatocyte cultures. Biochemical Pharmacology, 45(1), 13–19. https://doi.org/10.1016/0006-2952(93)90371-3

Nader K, Schafe GE, LeDoux JE (2000) Fear memories require protein synthesis in the amygdala for reconsolidation after retrieval. Nature 406:722–726.

Nader, K., and Einarsson, E. Ö. (2010). Memory reconsolidation: An update. Annals of the New York Academy of Sciences, 1191(1), 27–41. https://doi.org/10.1111/j.1749-6632.2010.05443.x

Nakagawa, T., Itoh, M., Ohta, K., Hayashi, Y., Hayakawa, M., Yamada, Y., Akanabe, H., Chikaishi, T., Nakagawa, K., Itoh, Y., Muro, T., Yanagida, D., Nakabayashi, R., Mori, T., Saito, K., Ohzawa, K., Suzuki, C., Li, S., Ueda, M., … Inuzuka, T. (2016). Improvement of memory recall by quercetin in rodent contextual fear conditioning and human early-stage Alzheimer’s disease patients. NeuroReport, 27(9), 671–676. https://doi.org/10.1097/WNR.0000000000000594

Orr, M. V., and Lukowiak, K. (2008). Electrophysiological and Behavioural Evidence Demonstrating That Predator Detection Alters Adaptive Behaviours in the Snail Lymnaea. Journal of Neuroscience, 28(11), 2726–2734. https://doi.org/10.1523/JNEUROSCI.5132-07.2008

Parvez, K. (2005). Boosting intermediate-term into long-term memory. Journal of Experimental Biology, 208(8), 1525–1536. https://doi.org/10.1242/jeb.01545

Parvez, Kashif, Moisseev, V., and Lukowiak, K. (2006). A context-specific single contingent-reinforcing stimulus boosts intermediate-term memory into long-term memory. European Journal of Neuroscience, 24(2), 606–616. https://doi.org/10.1111/j.1460-9568.2006.04952.x

Philips, G. T., Tzvetkova, E. I., Marinesco, S., and Carew, T. J. (2006). Latent memory for sensitization in Aplysia. Learning and Memory, 13(2), 224–229. https://doi.org/10.1101/lm.111506

Reznichenko, L., Amit, T., Zheng, H., Avramovich-Tirosh, Y., Youdim, M. B. H., and Mandel, S. (2006). Reduction of iron-regulated amyloid precursor protein and β-amyloid peptide by (-)-epigallocatechin-3-gallate in cell cultures: Implications for iron chelation in Alzheimer’s disease: Iron-mediated reduction of amyloid precursor protein. Journal of Neurochemistry, 97(2), 527–536. https://doi.org/10.1111/j.1471-4159.2006.03770.x

Ribeiro, M. J., Serfozo, Z., Papp, A., Kemenes, I., O’Shea, M., Yin, J. C. P., Benjamin, P. R., and Kemenes, G. (2003). Cyclic AMP response element-binding (CREB)-like proteins in a molluscan brain: Cellular localization and learning-induced phosphorylation. The European Journal of Neuroscience, 18(5), 1223–1234. https://doi.org/10.1046/j.1460-9568.2003.02856.x

Rivi, V., Benatti, C., Colliva, C., Radighieri, G., Brunello, N., Tascedda, F., and Blom, J. M. C. (2020). Lymnaea stagnalis as model for translational neuroscience research: From pond to bench. Neuroscience and Biobehavioural Reviews, 108, 602–616. https://doi.org/10.1016/j.neubiorev.2019.11.020

Rivi, V., Benatti, C., Lukowiak, K., Colliva, C., Alboni, S., Tascedda, F., and Blom, M. C. (2021b). What can we teach *Lymnaea* and what can *Lymnaea* teach us? Biological Reviews, brv.12716. https://doi.org/10.1111/brv.12716

Rivi, V., Batabyal, A., Benatti, C., Blom, J. M. C., and Lukowiak, K. (2021a). To eat or not to eat: A Garcia effect in pond snails (*Lymnaea stagnalis*). Journal of Comparative Physiology A. Accepted

Robak, J., and Gryglewski, R. J. (1988). Flavonoids are scavengers of superoxide anions. Biochemical Pharmacology, 37(5), 837–841. https://doi.org/10.1016/0006-2952(88)90169-4

Rosenzweig, M. R., Bennett, E. L., Colombo, P. J., Lee, D. W., and Serrano, P. A. (1993). Short-term, intermediate-term, and long-term memories. Behavioural Brain Research, 57(2), 193–198. https://doi.org/10.1016/0166-4328(93)90135-D

Sadamoto, H., Kitahashi, T., Fujito, Y., and Ito, E. (2010). Learning-dependent gene expression of CREB1 isoforms in the molluscan brain. Frontiers in Behavioural Neuroscience, *4*. https://doi.org/10.3389/fnbeh.2010.00025

Sadamoto, H., Sato, H., Kobayashi, S., Murakami, J., Aonuma, H., Ando, H., Fujito, Y., Hamano, K., Awaji, M., Lukowiak, K., Urano, A., and Ito, E. (2004). CREB in the Pond Snail Lymnaea stagnalis: Cloning, Gene Expression and Function in Identifiable Neurons of the Central Nervous System. Journal of Neurobiology, 58, 455–466. https://doi.org/10.1002/neu.10296

Sangha, S., Scheibenstock, A. McComb, C. and Lukowiak, K. (2003a). Intermediate and long-term memories of associative learning are differentially affected by transcription versus translation blockers in Lymnaea. Journal of Experimental Biology, 206(10), 1605–1613. https://doi.org/10.1242/jeb.00301

Sangha, S., Morrow, R., Smyth, K., Cooke, R., and Lukowiak, K. (2003b). Cooling blocks ITM and LTM formation and preserves memory. Neurobiology of Learning and Memory, 80(2), 130–139. https://doi.org/10.1016/S1074-7427(03)00065-0

Sangha, S., Scheibenstock, A., and Lukowiak, K. (2003c). Reconsolidation of a Long-Term Memory in *Lymnaea* Requires New Protein and RNA Synthesis and the Soma of Right Pedal Dorsal 1. The Journal of Neuroscience, 23(22), 8034–8040. https://doi.org/10.1523/JNEUROSCI.23-22-08034.2003

Sangha, S., Scheibenstock, A., Martens, K., Varshney, N., Cooke, R., and Lukowiak, K. (2005). Impairing Forgetting by Preventing New Learning and Memory. Behavioural Neuroscience, 119(3), 787–796. https://doi.org/10.1037/0735-7044.119.3.787

Scheibenstock, A., Krygier, D., Haque, Z., Syed, N., and Lukowiak, K. (2002). The Soma of RPeD1 Must Be Present for Long-Term Memory Formation of Associative Learning in *Lymnaea*. Journal of Neurophysiology, 88(4), 1584–1591. https://doi.org/10.1152/jn.2002.88.4.1584

Schroeter, H., Bahia, P., Spencer, J. P. E., Sheppard, O., Rattray, M., Cadenas, E., Rice-Evans, C., and Williams, R. J. (2007). (-)Epicatechin stimulates ERK-dependent cyclic AMP response element activity and up-regulates GluR2 in cortical neurons. Journal of Neurochemistry, 101(6), 1596–1606. https://doi.org/10.1111/j.1471-4159.2006.04434.x

Semon R. (1921) The mneme. London: Allen, Unwin.

Silva, A. J., Kogan, J. H., Frankland, P. W., and Kida, S. (1998). CREB and memory. Annual Review of Neuroscience, 21, 127–148. https://doi.org/10.1146/annurev.neuro.21.1.127

Socci, V., Tempesta, D., Desideri, G., De Gennaro, L., and Ferrara, M. (2017). Enhancing Human Cognition with Cocoa Flavonoids. Frontiers in Nutrition, 4, 19. https://doi.org/10.3389/fnut.2017.00019

Spencer, J. P. E. (2007). The interactions of flavonoids within neuronal signalling pathways. Genes and Nutrition, 2(3), 257–273. https://doi.org/10.1007/s12263-007-0056-z

Spencer, J. P. E. (2008). Flavonoids: Modulators of brain function? British Journal of Nutrition, 99(E-S1), ES60–ES77. https://doi.org/10.1017/S0007114508965776

Spencer, J. P. E., Rice-Evans, C., and Williams, R. J. (2003). Modulation of Pro-survival Akt/Protein Kinase B and ERK1/2 Signaling Cascades by Quercetin and Its in Vivo Metabolites Underlie Their Action on Neuronal Viability. Journal of Biological Chemistry, 278(37), 34783–34793. https://doi.org/10.1074/jbc.M305063200

Squire, L. R., Genzel, L., Wixted, J. T., and Morris, R. G. (2015). Memory Consolidation. Cold Spring Harbor Perspectives in Biology, 7(8). https://doi.org/10.1101/cshperspect.a021766

Stevens, C. F. (1994). CREB and memory consolidation. Neuron, 13(4), 769–770. https://doi.org/10.1016/0896-6273(94)90244-5

Sunada, H., Totani, Y., Nakamura, R., Sakakibara, M., Lukowiak, K., and Ito, E. (2017). Two Strains of Lymnaea stagnalis and the Progeny from Their Mating Display Differential Memory-Forming Ability on Associative Learning Tasks. Frontiers in Behavioural Neuroscience, *11*. https://doi.org/10.3389/fnbeh.2017.00161

Suzuki, A., Fukushima, H., Mukawa, T., Toyoda, H., Wu, L.-J., Zhao, M.-G., Xu, H., Shang, Y., Endoh, K., Iwamoto, T., Mamiya, N., Okano, E., Hasegawa, S., Mercaldo, V., Zhang, Y., Maeda, R., Ohta, M., Josselyn, S. A., Zhuo, M., and Kida, S. (2011). Upregulation of CREB-mediated transcription enhances both short- and long-term memory. The Journal of Neuroscience: The Official Journal of the Society for Neuroscience, 31(24), 8786–8802. https://doi.org/10.1523/JNEUROSCI.3257-10.2011

Swinton, E., de Freitas, E., Swinton, C., Shymansky, T., Hiles, E., Zhang, J., Rothwell, C., and Lukowiak, K. (2018). Green tea and cocoa enhance cognition in *Lymnaea*. Communicative and Integrative Biology, 11(1), e1434390. https://doi.org/10.1080/19420889.2018.1434390

Tronson, N. C., and Taylor, J. R. (2007). Molecular mechanisms of memory reconsolidation. Nature Reviews Neuroscience, 8(4), 262–275. https://doi.org/10.1038/nrn2090

Tully, T., Bourtchouladze, R., Scott, R., and Tallman, J. (2003). Targeting the CREB pathway for memory enhancers. Nature Reviews. Drug Discovery, 2(4), 267–277. https://doi.org/10.1038/nrd1061

Vauzour, D., Vafeiadou, K., Rice-Evans, C., Williams, R. J., and Spencer, J. P. E. (2007). Activation of pro-survival Akt and ERK1/2 signalling pathways underlie the anti-apoptotic effects of flavanones in cortical neurons. Journal of Neurochemistry, 103(4), 1355–1367. https://doi.org/10.1111/j.1471-4159.2007.04841.x

Widmer, Y. F., Bilican, A., Bruggmann, R., and Sprecher, S. G. (2018). Regulators of Long-Term Memory Revealed by Mushroom Body-Specific Gene Expression Profiling in Drosophila melanogaster. Genetics, 209(4), 1167–1181. https://doi.org/10.1534/genetics.118.301106

Williams, R. J., and Spencer, J. P. E. (2012). Flavonoids, cognition, and dementia: Actions, mechanisms, and potential therapeutic utility for Alzheimer disease. Free Radical Biology and Medicine, 52(1), 35–45. https://doi.org/10.1016/j.freeradbiomed.2011.09.010

Wilson, A. E., and Ross, M. (2003). The identity function of autobiographical memory: Time is on our side. Memory (Hove, England), 11(2), 137–149. https://doi.org/10.1080/741938210

Yevchak, A. M., Loeb, S. J., and Fick, D. M. (2008). Promoting Cognitive Health and Vitality: A Review of Clinical Implications. Geriatric Nursing, 29(5), 302–310. https://doi.org/10.1016/j.gerinurse.2007.10.017

Yin, J. C., and Tully, T. (1996). CREB and the formation of long-term memory. Current Opinion in Neurobiology, 6(2), 264–268. https://doi.org/10.1016/S0959-4388(96)80082-1

Yoshida, M., Sakai, T., Hosokawa, N., Marui, N., Matsumoto, K., Fujioka, A., Nishino, H., and Aoike, A. (1990). The effect of quercetin on cell cycle progression and growth of human gastric cancer cells. FEBS Letters, 260(1), 10–13. https://doi.org/10.1016/0014-5793(90)80053-l

Zlotnik, G., and Vansintjan, A. (2019). Memory: An Extended Definition. Frontiers in Psychology, 10. https://doi.org/10.3389/fpsyg.2019.02523

